# Prostate-derived circulating microRNAs add prognostic value to prostate cancer risk calculators

**DOI:** 10.1101/2023.05.10.540236

**Authors:** Morgan L. Zenner, Brenna Kirkpatrick, Trevor R. Leonardo, Michael J. Schlicht, Alejandra Cavazos Saldana, Candice Loitz, Klara Valyi-Nagy, Mark Maienschein-Cline, Peter H. Gann, Michael Abern, Larisa Nonn

**Author notes:** Correspondence: Larisa Nonn, PhD Professor of Pathology 840 S. Wood St. Chicago, IL 60612. The authors have declared that no conflict of interest exists.

## Abstract

Prostate cancer is the second leading cause of malignancy-related deaths among American men. Active surveillance is a safe option for many men with less aggressive disease, yet definitively determining low-risk cancer is challenging with biopsy alone. Herein, we sought to identify prostate-derived microRNAs in patient sera and serum extracellular vesicles, and determine if those microRNAs improve upon the current clinical risk calculators for prostate cancer prognosis before and after biopsy. Prostate-derived intracellular and extracellular vesicle-contained microRNAs were identified by small RNA sequencing of prostate cancer patient explants and primary cells. Abundant microRNAs were included in a custom microRNA PCR panel that was queried in whole serum and serum extracellular vesicles from a diverse cohort of men diagnosed with prostate cancer. The levels of these circulating microRNAs significantly differed between indolent and aggressive disease and improved the area under the curve for pretreatment nomograms of prostate cancer disease risk. The microRNAs within the extracellular vesicles had improved prognostic value compared to the microRNAs in the whole serum. In summary, quantifying microRNAs circulating in extracellular vesicles is a clinically feasible assay that may provide additional information for assessing prostate cancer risk stratification.

## Introduction

Prostate cancer (PCa) is the most diagnosed cancer and the second most common cause of death due to malignancy among men in the United States (US), with one in eight men in the US being diagnosed during their lifetime^1^. Because many men with PCa will die from other causes, a current challenge is determining which patients will develop aggressive, metastatic PCa, and which patients will have indolent, localized PCa^2^. The five-year relative survival rate drops from 100% in patients with localized PCa to 32.3% in patients with metastatic disease^3^, lending urgency to developing improved prognostic strategies.

PCa is diagnosed by a multiple-core prostate biopsy with aggressiveness determined by the Gleason grading system which includes a primary and secondary grade, ranging from 3 to 5, with 5 representing least differentiated anaplastic cells^4^. The grades are classified into a Gleason Grade Group (GG) ranging from 1-5, with GG1 as the lowest grade PCa (Gleason 3+3) and GG5 as the highest grade PCa (Gleason 4+5, 5+4, or 5+5)^5^. The challenge with biopsies is that even with MRI guidance, they are a somewhat random sampling of the peripheral zone, which can lead to false-negative results. Thus, a low Gleason grade on biopsy tissue does not rule out the presence of a high-grade tumor that was not biopsied^6^.

For PCa prognosis after biopsy, urologists combine GG with other clinical parameters to stratify patients into risk groups^7^. These risk groups predict the risk of biochemical recurrence (BCR), a rise in serum prostate specific antigen (PSA) after primary treatment, which is indicative of metastases. Risk group and life expectancy are used to guide treatment recommendations in a shared decision process with the patient. Treatment options for localized PCa include active surveillance, radical prostatectomy (RP), radiation therapy (RT), cryotherapy, and high-intensity– focused ultrasound^7^. Compared with active surveillance, RP and RT decrease PCa metastasis and mortality rate ^8–10^, but are associated with adverse side effects such as urinary incontinence and erectile dysfunction^11^. Moreover, biopsy GG does not always align with GG of the RP as 30-45% of patients have upgraded and 10% have downgraded GG at RP^6^, contributing to the difficulties faced by patients and clinicians when making PCa treatment choices.

Multiple nomograms and tissue-based tests have been developed to provide personalized risk predictions at various stages of diagnosis ^7, 12–14^. The Prostate Biopsy Collaborative Group (PBCG) risk calculator is assessed before a biopsy to predict the likelihood of a positive biopsy^12^. Following a PCa-positive biopsy, the Cancer of the Prostate Risk Assessment (CAPRA) nomogram, is used to aid in treatment decisions by predicting adverse pathology (AP) of the prostate^14, 15^. These nomograms take into account family history, serum PSA and other clinical parameters. Multiple biopsy tissue-based prognostic tests have been developed to further guide treatment decisions ^6, 16^, yet Salami et al. found significant differences between patient-matched GG1 and GG4 prostate cores when comparing three tissue-based tests, indicating that transcription profiles from lower-grade biopsy cores do not capture the profiles from higher-grade cores^17^.

MicroRNAs (miRs) are highly stable in circulation^18^ and are promising prognostic biomarkers for PCa in prostate tissues and multiple biological fluids as shown by our group and others^19–21^. Our group previously examined 14 serum miRs in a cohort of 150 men with high-grade PCa, low-grade PCa or benign prostate hyperplasia and determined they were predictive of indolent PCa or a cancer-free status with a negative predictive value of 100% and positive predictive value of 58.8%^21^. Circulating miRs are often enclosed within extracellular vesicles (EVs)^19^, a heterogeneous group of vesicles with lipid bilayer membranes which includes exosomes^22^. EVs are released from cells and have functions in normal physiology and in various disease processes ^23, 24^. Due to the stability and functionality of miRs in EVs, they have gained substantial attention as liquid biomarkers and mediators of various cancers, including PCa ^25, 26^.

Herein, we sought to first identify prostate-derived miRs in EVs from patient cells and tissues and then to determine the prognostic ability of serum and serum EV miRs in a diverse cohort of 203 treatment-naïve patients with biopsy-confirmed PCa. The primary endpoint was assessing the ability of circulating miRs to add prognostic value to the current clinical standard, the CAPRA nomogram, for determining adverse pathology status at RP. The secondary endpoint involved determining if the circulating miRs added to the prognostic value of the PBCG nomogram for discriminating low-grade from high-grade PCa at biopsy.

## Methods

### Patient-derived prostate tissue and cell cultures

Patient prostate cells and explants were derived from patient radical prostatectomy specimens via informed consent prior to surgery (UIC Internal Review Board-approved protocol #2006-0679). The tissue was either sliced for explant culture or dissociated for cell propagation as previously described by our lab ^27^. Briefly, dissociated cells were grown in Prostate Cell Growth Media (PrEBM) (Lonza, Basel, Switzerland) or MCDB105, for prostate epithelial (PrE) and stromal (PrS) cells, respectively. Cell type authentication was by RT-qPCR; epithelial-specific markers were keratin 5 (KRT5) (F: CCATATCCAGAGGAAACACTGC, R: ATCGCCACTTACCGCAAGC) and keratin 18 (KRT18) (F: CACAGTCTGCTGAGGTTGGA, R: CAAGCTGGCCTTCAGATTTC), and the stromal-specific markers were vimentin (VIM) (F: CGAAAACACCCTGCAATCTT, R: TCCTGGATTTCCTCTTCGTG) and Androgen Receptor (AR) (F: CCAGGGACCATGTTTTGCC, R: CGAAGACGACAAGATGGACAA). For explant cultures, 300 μm tissue slices were generated using a Tissue Slicer (Model MD6000, Alabama R&D, Munford, AL, USA) and cultured on titanium grids that were constantly rotated in and out of PrEBM containing 50 nM R1881.

### EV isolation from cells

Prior to EV isolation, cells were cultured in EV-free medium (bovine supplement components omitted) for 48h. PrE and PrS EVs were isolated from conditioned medium using differential ultracentrifugation; 300 x g for 10 minutes (min), supernatant centrifuged at 2,000 x g for 20 min, supernatant was ultracentrifuged at 10,000 x g for 30 min, supernatant ultracentrifuged at 100,000 x g for 16h to pellet EVs. The EV pellet was washed in 1X phosphate-buffered saline (PBS)(no magnesium or calcium) (Corning Inc., Corning, NY, USA), ultracentrifuged for 70 min at 100,000 x g and resuspended in 100 µL 1X PBS. All centrifugation and ultracentrifugation steps were performed at 4°C ^28^.

### EV isolation from tissues

EVs were isolated from 2.5 mL of 48 h conditioned media from tissue slice explants using ExoQuick-TC (System Biosciences, Palo Alto, CA, USA) per the manufacturer’s protocol. The EV pellet was resuspended in 100 µL 1X PBS (no magnesium, no calcium) or purified via a bipartite resin column according to the ExoQuick-TC ULTRA protocol.

### EV isolation from serum

Serum was collected in BD gold-top (serum-separator) vacutainers, incubated for 30 min at room temperature (RT), centrifuged at 1,315 x g for 10 min at 4°C. RNase inhibitors were added to the separated serum; 40 units/µL RNaseOUT (Life Technologies, Carlsbad, CA, USA) and 20 units/µL SUPERase-IN (Life Technologies, Carlsbad, CA, USA). EVs were isolated from 500 µL fresh serum using the ExoQuick protocol (System Biosciences, Palo Alto, CA, USA). Remaining serum aliquots were snap-frozen and stored at -80°C.

### Nanoparticle Tracking Analysis (NTA)

EV pellets were run on the NanoSight NS300 according to the instrument protocol (MAN0541-01-EN-00, 2017). For each sample, three 30-second videos were captured using a 488 nm laser and a syringe pump speed of 100 (Malvern Instruments, United Kingdom) and automatically analyzed using the built-in software NTA 3.2 (Malvern Instruments, Westborough, MA, USA) using a detection threshold of 5. For the particle concentration and size analyses, particle counts were binned in 10 nm increments. The results from the control sample (PBS) were subtracted from each sample run to determine the concentration of EVs within each 10 nm size bin.

### Transmission Electron Microscopy (TEM)

EV sample in 10-15 µL 1X PBS was deposited onto a 300-mesh Formvar/Carbon-coated copper EM grid and fixed with 2% Uranyl acetate solution (heavy metal stain). The grid was dried and examined immediately via TEM. EV samples were examined with the JEOL JEM-1400F transmission electron microscope operating at 80 kV. Digital micrographs were acquired using an AMT NanoSprint1200-S camera and AMT software (version 7.01).

### CD63 ELISA

EV pellets were diluted and 45 µL was examined with the ExoELISA-ULTRA CD63 Kit according to the manufacturer’s instructions (System Biosciences, Palo Alto, CA, USA). The sample absorbance minus the blank wells was used to determine concentrations based on the standard curve included in the kit.

### Western Blot

The EV pellet was resuspended in RIPA buffer with protease/phosphatase inhibitor (Cell Signaling Technology, Danvers, MA, USA). Protein (20 µg) was loaded in Laemmli buffer, run on a 4-12% Bis-Tris electrophoresis gel (Life Technologies, Carlsbad, CA, USA), and transferred to polyvinylidene difluoride (PVDF) membrane (Merck Millipore, Carrigtwohill, CO, USA). Blot was blocked with 5% dry milk in Tris-buffered saline plus 0.05% Tween (TBST) and incubated overnight at 4°C with 1:1000 primary antibody (CD63 and CD81, rabbit anti-human, EXOAB-CD63A-1/ EXOAB-CD81A-1, System Biosciences, Palo Alto, CA, USA) in TBST. Membranes were washed, incubated in 1:20,000 secondary antibody (Goat Anti-Rabbit HRP) and visualized with SuperSignal West Atto Ultimate Sensitivity Chemiluminescent Substrate (Thermo Fisher Scientific, Waltham, MA, USA) on the iBright 1500 (Thermo Fisher Scientific, Waltham, MA, USA).

### MicroRNA RNAseq library preparation and sequencing

PrE and PrS cells were cultured to 70% confluency, conditioned media was collected, and EVs were isolated by differential ultracentrifugation, as described above. PrE and PrS cells were directly lysed, and RNA was collected using QIAzol (Qiagen, Germany). Tissue slices were homogenized in QIAzol using the BeadRuptor 4 (OMNI International, Bedford, NH, USA). RNA was isolated from all intracellular samples with the miRNeasy Mini Kit (Qiagen, Germany) and from EVs using the Serum/Plasma miRNeasy Kit (Qiagen, Germany). MiR quantification from all samples was completed with microRNA Qubit (Thermo Fisher Scientific, Waltham, MA, USA).

MiR libraries were generated from 100 -150 ng of cell using the QIAseq miRNA Library Kit according to the protocol (Qiagen, Germany). Library quality control and concentrations were determined using a High Sensitivity D1000 ScreenTape with TapeStation Analysis Software A.02.02 (Agilent Technologies, Santa Clara, CA, USA) Libraries (10 µM) were submitted to the Roy J. Carver Biotechnology Center at the University of Illinois at Urbana-Champaign, where they were titrated for equal loading using the MiSeq followed by sequencing on a HiSeq 4000 with 100 nucleotide forward single reads.

### MicroRNA differential expression analysis

Raw FASTQ sequencing reads were inputted into the GeneGlobe Data Analysis Center (Qiagen, Germany). Reads were trimmed using cutadapt ^29^, which removed the 3’ adapter and low-quality bases. MiR sequences less than 16 base pairs (bp) and UMI sequences less than 10 bp were removed. All samples passed the initial quality control checkpoints in the GeneGlobe Data Analysis Center. Reads were then mapped to miRBase (version 22) using bowtie ^30^, which allows up to two mismatches. Raw counts were generated by calculating the number of UMIs mapped to each miR sequence. Downstream analyses were performed using R version 3.4.1 ^31^.

PrE, PrS, tissue slice (TSC), PrE EV, PrS EV, and TSC EV samples were initially grouped together with Serum EV samples, and piwi-interacting RNA (piRNA) were removed. Next, miRs were filtered such that only miRs with at least five raw reads in at least 50% of all samples were retained, leaving 2114 of the original 2523 miRs. The data was then separated into the following four datasets: 1) PrE EV, PrS EV, TSC EV, Serum EV; 2) PrE EV, PrS EV, TSC EV; 3) Serum EV; and 4) PrE, PrS, TSC. For each individual dataset, a DGEList object was created in EdgeR ^32^, normalization factors were calculated using the default trimmed mean of M-values (TMM) between each pair of samples, counts per million (CPM) function with or without log2 scaling was used to generate CPM expression levels of each miR, and files were saved. Log2 scaled CPM data were used to generate all heatmaps and principal component plots (PCA) using the heatmap3, ggplot2, and cowplot packages ^33, 34^. Differential expression analyses were performed using EdgeR. For each dataset, a DGEList object was created, normalization factors were calculated using TMM, and genewise statistical tests were conducted using the estimateDisp, glmQLFit, glmQLFTest, and topTags functions to identify genes differentially expressed between any of the groups within each dataset. For heatmaps, only differentially expressed genes with a statistically significant adjusted p-value ≤ 0.01 by Benjamini and Hochberg method were plotted^35, 36^.

### Prostate cancer patient cohort description

Serum was collected from two UIC Urology PCa patient cohorts; one cohort included patients on active surveillance as part of the Engaging Newly Diagnosed Men About Cancer Treatment Options (ENACT) trial (IRB protocol # 2015-1294), and the second cohort included patients who had chosen RP treatment for PCa (Prostate Cancer Blood Biorepository Resource [PCBBR], IRB protocol # 2017-0807). ENACT trial patients were 76 years or younger with a recent diagnosis of very low-, low-, or low-intermediate risk PCa. The only inclusion criterion for the PCBBR cohort was the decision to undergo RP. Due to the differences in inclusion criteria, the ENACT cohort consisted of lower-risk PCa patients (biopsy Gleason grade groups 1-2), and the PCBBR cohort consisted of relatively higher-risk PCa patients (biopsy Gleason grade groups 1-5). Age, PSA level, self-declared race, biopsy Gleason grade group, and pathological Gleason grade group data are included for each cohort. For the ENACT patient cohort, serum was collected at least eight weeks post-biopsy and prior to RP treatment (if the patient terminated active surveillance treatment). Serum was collected at least one month after biopsy but prior to RP treatment for the PCBBR cohort. Serum samples were immediately processed upon collection, separated into 500 µL aliquots, snap-frozen, and stored at -80°C. EVs were isolated immediately from serum and stored in liquid nitrogen.

### Custom reverse-transcription quantitative polymerase chain reaction (RT-qPCR) miR panel

Five µL RNA was reverse transcribed to cDNA using the miRCURY LNA RT Kit (Qiagen, Germany). MiRs (61 plus 3 controls) were quantified on a custom PCR plate using the miRCURY LNA miR SYBR Green PCR Handbook (Qiagen, Germany). PCR (40 cycles) was run according to the manufacturer’s instructions.

### Normalization and filtering of qPCR values

NormFinder normalization values were generated by Gene Globe. The normalization panel for the serum samples, as determined by NormFinder, included hsa-miR-103a-3p, hsa-miR-107, hsa-miR-24-3p, hsa-miR-30c-5p, hsa-miR-93-5p, hsa-miR-19b-3p, hsa-miR-222-3p, hsa-miR-27a-3p, hsa-miR-23a-3p, and hsa-let-7b-5p. The normalization panel for the serum EV samples, as determined by NormFinder, included hsa-miR-107, hsa-miR-24-3p, hsa-miR-30c-5p, hsa-miR-93-5p, hsa-let-7i-5p, hsa-miR-222-3p, hsa-miR-27a-3p, hsa-miR-23a-3p, hsa-miR-21-5p, and hsa-miR-27b-3p. Because each normalization factor was based on the average of 10 miRs, these miRs were included in the analysis for significance based on outcomes.

The call rate per sample was computed as the fraction of miRs with Ct lower than 33, and samples with a call rate <50% were removed from the dataset. Similarly, miRs with a call rate <60% were removed. Normalized delta-Ct values were computed and miRs with Ct ≥33 were set to the minimum normalized expression over all other samples for the same miR. MiRs from serum and extracellular vesicles (EVs) were processed independently in the identical manner.

### Statistics—Random Forest Model

Statistical analyses were completed at the UIC Research Resource Center Bioinformatics Core. The following binary outcomes were considered in preparing the models: (1) Biopsy Gleason grade group 1 (low-grade) and Gleason grade group ≥ 2 (high-grade) and (2) adverse pathology or non-adverse pathology at RP. For each binary outcome, samples with missing values were removed, and *t* tests were run for each miR using data from serum and EVs separately. MiRs with a p-value < 0.05 were retained as features in the random forest model. We also included clinical features for different models: PBCG high-grade, low-grade, and negative biopsy percentages were included independently or in combination with miRs for the prediction of Gleason grade group outcomes. CAPRA score was included independently or in combination with miRs for the prediction of adverse pathology.

For each outcome and feature dataset, we trained and tested the performance of the random forest model using the randomForest() function in the randomForest package in R^37^ with default parameters. Leave-one-out cross validation was performed, predicting the outcome of each sample based on a model trained on all other samples. ROC and AUC statistics were computed based on cross validation predictions using the roc() function in the pROC package in R (46). Confidence intervals of AUC statistics were computed via bootstrapping with 500 bootstrap replicates using the ci.auc() function. P-values comparing the pairwise performance of different models were computed using the Wilcoxon signed rank test between the predicted numerical values from each model.

### Study Approval

Patient specimens for this study were collected by three UIC IRB-approved protocols, one for prostate tissue/cells and two for sera, #2006-0679, #2015-1294, and #2017-0807.

### Data Availability

RNA sequencing and RT-qPCR files are available at GEO Accession numbers: GSE228062, GSE228371, GPL33285. R-code is available at https://codeocean.com/capsule/1582037/tree.

## RESULTS

### Distinct cell-type specific miR expression profiles for prostate-derived cells, tissues, and EVs

The prostate is composed of multiple cell types, with a fibromuscular stroma surrounding the glandular epithelium from which cancer arises. To identify prostate-derived miRs, the miR profiles of patient-derived prostate tissues, patient-derived primary prostate epithelial cells (PrE), patient-derived primary prostate stromal cells (PrS), and their EVs were characterized by small RNA next-generation sequencing (small RNAseq) (**Figure 1A**). Ex vivo prostate tissue slice cultures (TSC) (N=4) were used as the source of whole tissue miRs and TSC EVs. EVs were isolated from conditioned media from patient-derived PrE and PrS samples (N=4 patients). Cell type was verified by gene expression of known markers (**Supplementary Figure 1**).

**Figure 1.**
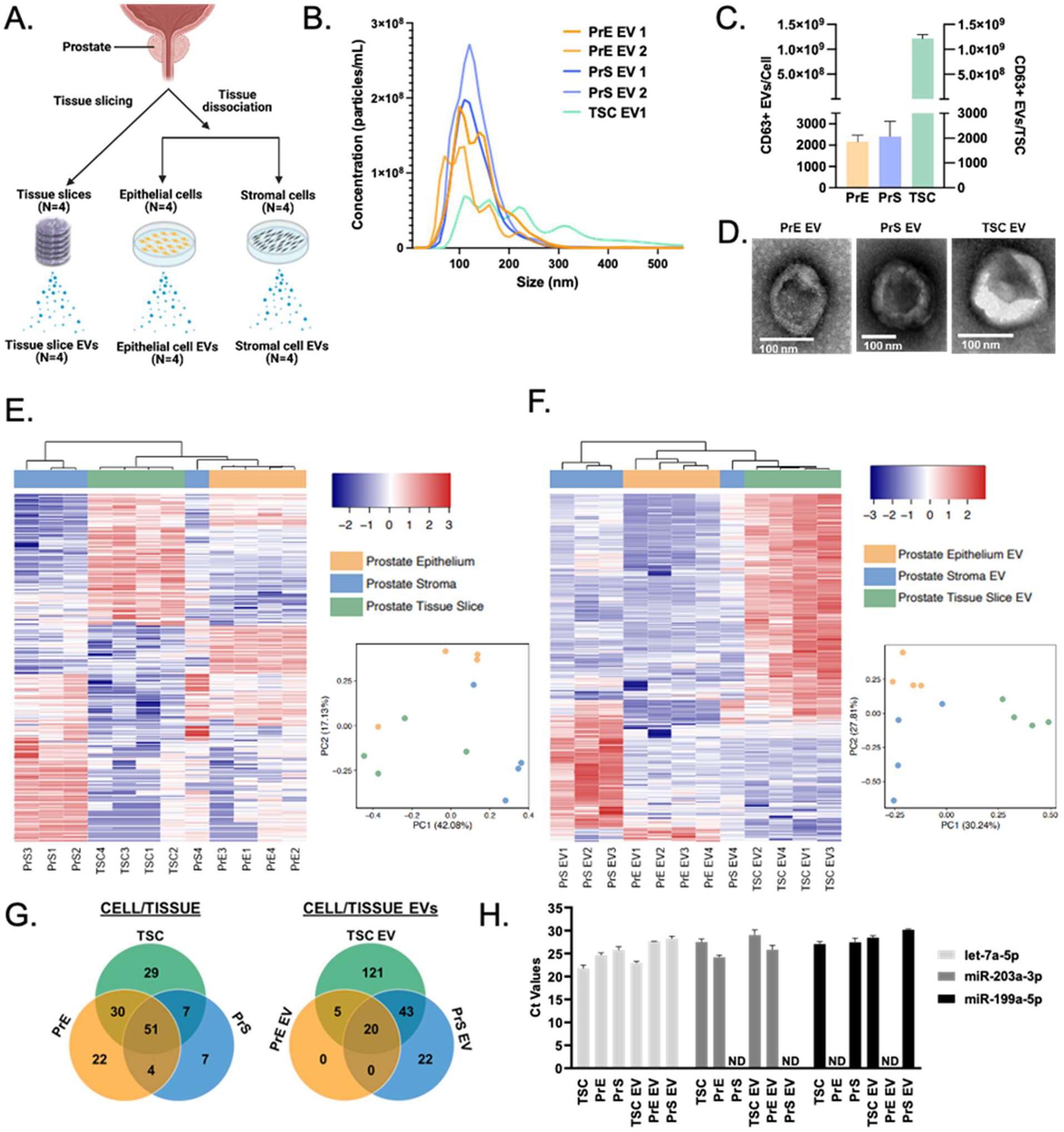
Characterization of prostate-derived extracellular vesicles and microRNA profiles of patient-derived prostate samples. **A)** Flow chart of PCa patient-derived samples included for small RNA NGS profiling (4 tissue slice explants (TSC), 4 epithelial cell cultures (PrE), 4 stromal cell cultures (PrS), and 4 EV samples from each sample type (TSC EV, PrE EV, and PrS EV). **B)** Nanoparticle particle tracking analysis of two PrE EV samples, two PrS EV samples, and one TSC EV sample. **C)** CD63 ELISA of PrE-, PrS-, and TSC-derived EVs. (N = 3 PrE, 3 PrS, 2 TSC; Error bars show SEM). **D)** Transmission electron microscopy (TEM) images of EVs derived from PrE, PrS, and TSC (size bars = 100 nm). **E, F)** PCA plots and unbiased hierarchical clustering of the miR profiles of (**E**) Patient-derived prostate epithelial cells, stromal cells, and tissue slice explants and (**F**) EVs isolated from patient-derived prostate epithelial cells, stromal cells, and tissue slice explants (N = 4 per sample type). Heatmaps show differentially expressed miRs with an adjusted p-value ≤ 0.01 using the Benjamini-Hochberg method. **G)** Venn diagram representation of TMM-normalized miRs with > 500 counts per million (cpm) within each sample type. **H)** RT-qPCR validation of cell/EV-specific miRs as determined by NGS miR profiles (N = 2 replicates per sample, Error bars = SD, ND = not detected).

TSC, PrE, and PrS EVs were characterized by Nanosight Tracking Analysis (NTA), CD63 ELISA, and transmission electron microscopy (TEM) (**Figure 1B-D**). The EVs varied in size between the samples, with PrE EVs showing one uniform peak, and PrS and TSC EVs showing more heterogeneity with multiple size peaks (**Figure 1B**). EVs were CD63-positive in all samples (**Figure 1C**), and TEM of TSC, PrE, and PrS EVs showed sizes of 50-200 nm and a classic cup-shaped morphology (**Figure 1D**).

Small RNA libraries were created and analyzed using EdgeR (38). Principal component analysis (PCA) showed that both intracellular and EV miRs clustered by sample type (**Figures 1E-F**). Of note, PrS4 and PrS EV4 did not cluster tightly with the other three PrS samples and clustered more closely with the PrE and PrE EV samples, respectively. One of the PrE samples clustered more tightly with the TSC samples.

Differential expression analysis in EdgeR identified common miRs and those with distinct expression in TSC, PrE, and PrS samples and EV samples. Unbiased hierarchical clustering heatmaps of miRs showed grouping by sample type (q ≤ 0.01, **Figure 1E-F, Supplementary Tables 1-2**), which was similar to the PCA. To determine unique and common miRs, miRs were considered “detectable” with at least 500 normalized counts per million (cpm). For intracellular miRs, TSC expressed the most miRs and the majority of cell-type specific miRs were also detected in the TSCs (**Figure 1G and Supplementary Table 3**). Similarly, the TSC EVs had the most detectable miRs (**Figure 1G** and **Supplementary Table 4).** RT-qPCR was used to validate a few of the cell type-specific miR expression profiles observed by small RNA-seq. Hsa-let-7a-5p was robustly expressed in all sample types (PrE, PrS, TSC, and EVs), supporting the small RNAseq (**Figure 1H**). Hsa-miR-203a-3p was confirmed to be specific to PrE/PrE EVs and TSC/TSC EV (**Figure 1H**) whereas hsa-miR-199a-5p was specific to PrS/PrS EV and TSC/TSC EV (**Figure 1H**). MiR profiling showed that the stroma and epithelium have distinct EV miR populations and that almost all stromal and epithelial EV miRs were present in the EVs from ex vivo prostate tissue slices, which contain both epithelial and stromal components.

### Prostatic miRs within serum extracellular vesicles as biomarkers in prostate cancer patients

EVs were isolated freshly from patient sera. NTA and TEM showed the serum EV size 50-300 nm (**Figure 2A-B**). Serum EVs expressed CD63 and CD81 but not GRP78 (intracellular protein) by western blot (**Figure 2C**). Serum miRs are known to be exceptionally stable (22), but the stability of serum miRs packaged in EVs has not been well described. To mimic potential delays and storage conditions that could occur in the clinic, EVs were immediately isolated or stored at 4°C, -20°C, or -80°C for seven days. CD63 levels remained unchanged after a seven-day delay in EV isolation (**Figure 2D**) and the expression levels of known serum miRs remained consistent in all conditions (**Supplementary Figure 2).**

**Figure 2.**
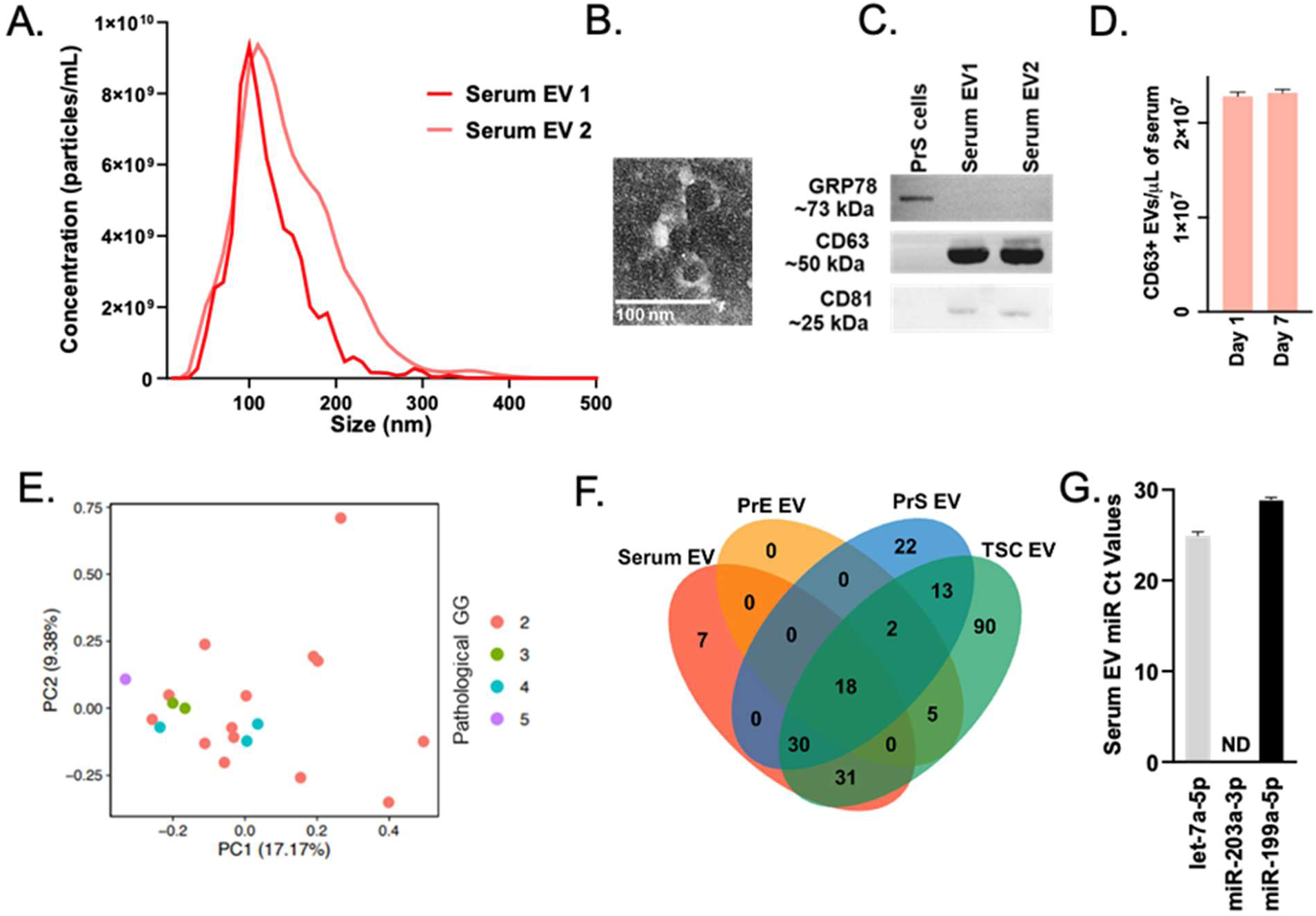
Characterization of patient-derived pretreatment serum extracellular vesicles and their microRNA profiles. **A)** Nanoparticle tracking analysis of N = 2 patient-derived serum EV samples. **B)** Transmission electron microscopy image of patient-derived serum EVs (size bar = 100 nm). **C)** Western blot of EV-specific markers (CD63 and CD81) and intracellular protein control (GRP78) of two patient-derived serum EV samples and one patient-derived stromal cell control. **D)** CD63 ELISA of patient-derived serum EV samples stored at 4, -20, and -80°C for 1 and 7 days. (N = 3 samples per day, error bars = SD). **E)** PCA plot of the miR profiles of patient-derived pretreatment serum EVs (N = 20 patients). Data are grouped by pathological Gleason Grade group (Pathological GG). **F)** Venn diagram representation of TMM-normalized miRs with > 500 counts per million (cpm) within EVs isolated from patient-derived serum, epithelial cells, stromal cells, and tissue slice explants. **G)** RT-qPCR validation of serum EV miRs as determined by NGS miR profiles (N = 2 replicates per sample, Error bars = SD, ND = not detected).

To identify potential biomarkers for PCa, serum EV miR expression profiles from PCa patients (N=20) were examined by small RNAseq. PCA showed no clustering by the pathological Grade Group, but only six patients were higher than GG2 (**Figure 2E**). MiRs from the serum EV miRs were compared to the profiles from the TSC EVs, PrE EVs, and PrS EVs (**Figure 2F**) to identify potential prostate-derived miRs in the serum. There were 18 detectable miRs in common between serum EV, TSC EV, PrE EV, and PrS EV (**Figure 2F**). By RT-qPCR, hsa-let-7a-5p and hsa-miR-199a-5p were present in serum EVs and hsa-miR-203a-3p was not detected, validating the small RNAseq findings (**Figure 2G).**

### Whole serum and serum EV microRNAs add prognostic value to pre-treatment clinical nomograms

A prospective study was designed to examine the prognostic ability of post-biopsy serum miRs in UIC PCa patients (N=203) (**Figure 3A and Table 1**). Serum was prospectively collected from two PCa patient cohorts with same-day EV isolation. As the adverse pathology classification of RP resections is a major predictor of disease recurrence after RP, the primary goal of this study was to identify miRs in presurgical serum that are prognostic for RP tumor pathology. Hence, the custom miR panel was run on the serum (N=81) and serum EVs (N=62) from patients who later underwent RP. Adverse pathology is defined as Gleason Grade Group 4 or 5 and/or extraprostatic disease at RP and is a hallmark of aggressive disease that is likely to progress ^38^.

**Figure 3.**
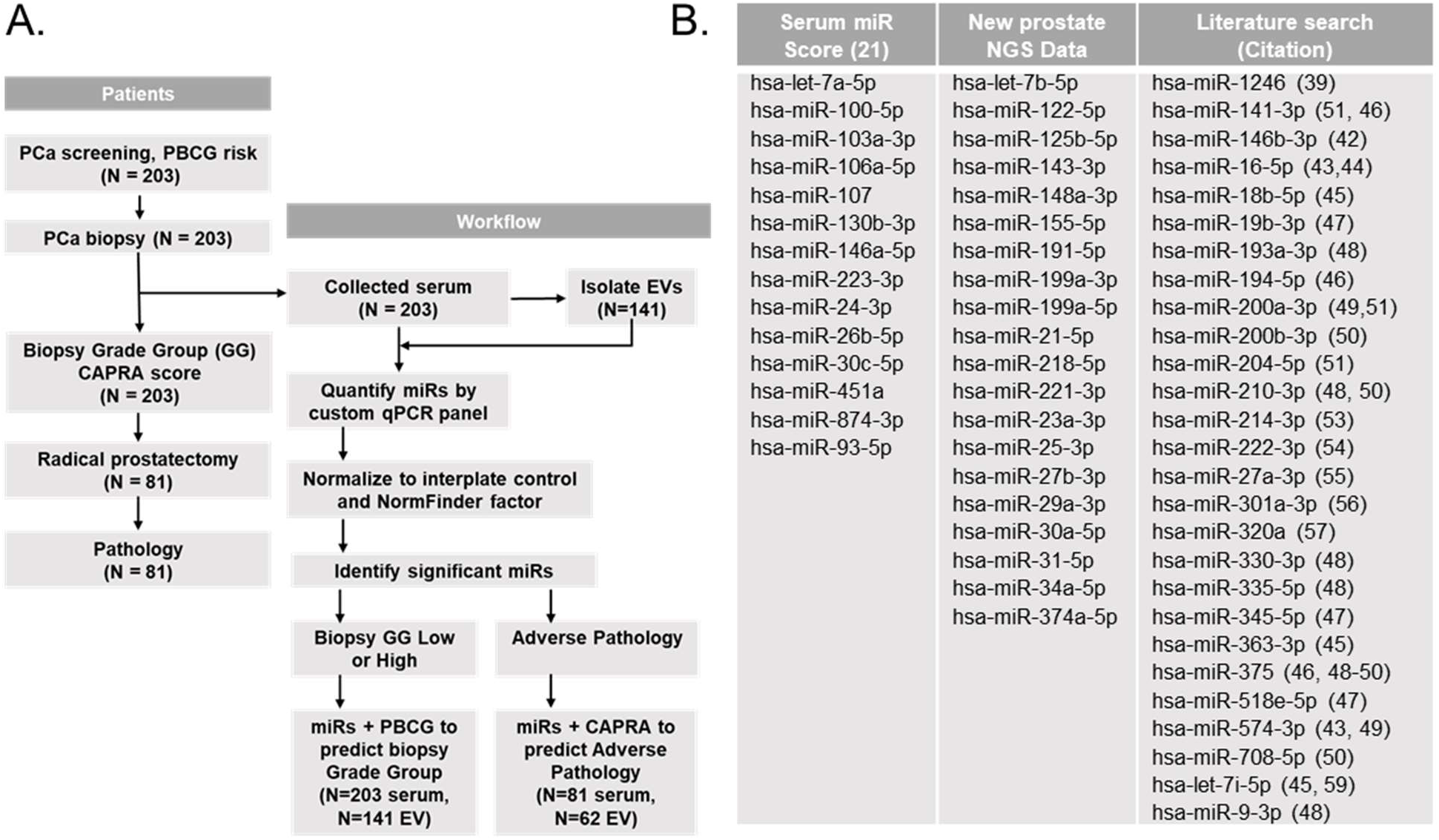
Patient cohort and study design. **A)** Study design flow chart for determining serum and serum EV miRs that are predictive of prostate cancer outcomes. **B)** MiRs in the panel and how they were selected, either from our prior study, the new NGS data, or a literature search.

**Table 1.** Patient cohort characteristics.

| **CHARACTERISTICS** |  | **ENACT** | **PCBBR** |
| --- | --- | --- | --- |
| **AGE (Median with Range)** |  | 62 (47-76) | 62 (45-77) |
| **PSA ng/mL (Median with Range)** |  | 5.58 (0.47-14.17) | 7.05 (3-735) |
| **RACE**<br>**N (%)** | **African American** | 95 (67.4) | 33 (53.2) |
|  | **Caucasian** | 25 (17.7) | 15 (24.2) |
|  | **Hispanic** | 19 (13.5) | 10 (16.1) |
|  | **Asian** | 2 (1.4) | 0 (0) |
|  | **Multiple Races/Unknown** | 0 (0) | 4 (6.5) |
| **BIOPSY GLEASON**<br>**GRADE GROUP**<br>**N (%)** | **1** | 119 (84.4) | 7 (11.3) |
|  | **2** | 22 (15.6) | 34 (54.8) |
|  | **3** | 0 (0) | 11 (17.7) |
|  | **4** | 0 (0) | 7 (11.3) |
|  | **5** | 0 (0) | 3 (4.8) |
| **BIOPSY TO RP**<br>**GLEASON GRADE**<br>**COMPARISON\***<br>**N (%)** | **Upgraded** | 10 (40.0) | 17 (29.3) |
|  | **Downgraded** | 0 (0) | 13 (22.4) |
|  | **Consistent** | 15 (60.0) | 28 (48.3) |
| **ADVERSE**<br>**PATHOLOGY**<br>**N (%)** | **Yes** | 8 (32.0) | 36 (58.1) |
|  | **No** | 17 (68.0) | 20 (32.2) |
|  | **Unknown** | 0 (0) | 6 (9.7) |

\*Gleason Grade groups (GG) were compared at biopsy and at RP; patients with the same GG were qualified as "Consistent", patients with higher GG at RP compared to biopsy were qualified as "Upgraded", and patients with lower GG at RP compared to biopsy were qualified as "Downgraded".

Based on the RNA-seq data and current literature, we created a custom qPCR panel to quantify 61 miRs in whole serum and serum EV samples. The miRs included in the panel were the 14 miRs from our previously reported Serum miR Score^21^, 20 miRs that were highly expressed in prostate cell or tissue EVs and serum EVs by RNAseq analysis, and 27 miRs that were consistently reported in the literature as prognostic serum biomarkers for PCa^18,39–59^ (**Figure 3B**). MiRs significantly different between patients with no adverse pathology and patients with adverse pathology in serum and serum EV samples were determined and included in the random forest model analyses.

CAPRA score, a clinical post-biopsy nomogram, was used for comparison ^60^. The outputs for CAPRA are low risk (score 0-2), intermediate risk (score 3-5), and high risk (score 6-10) ^61^. Random forest models showed that serum miRs and serum EV miRs added to the prognostic power of CAPRA for adverse pathology. Seven serum miRs significantly improved upon the CAPRA score AUC for adverse pathology, 0.675 (95% CI:0.547-0.799), compared to the CAPRA score alone, with an AUC of 0.516 (95% CI:0.379-0.643, p = 0.010) (**Figure 4A,C** and **Supplementary Table 5**). 19 serum EV miRs were significantly different in patients with adverse pathology and improved upon the CAPRA model (**Figure 4B,C** and **Supplementary Table 5**). Adding serum EV miRs to the CAPRA model increased the AUC to 0.739 (95% CI:0.580-0.900) (p = 0.010). Some, but not all, of the significant miRs differed between serum and serum EVs.

**Figure 4.**
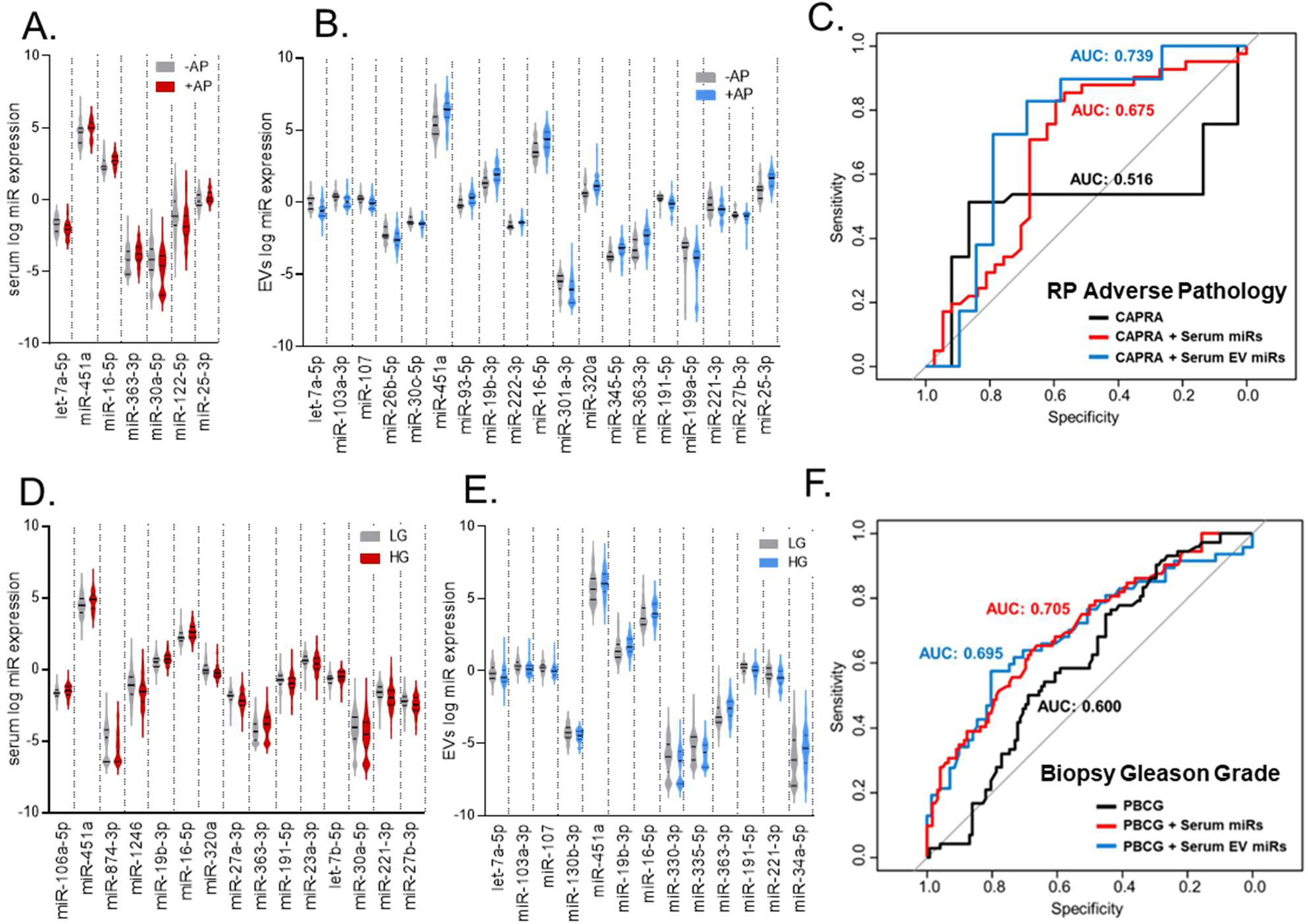
Circulating serum and serum EV microRNAs significantly improve AUCs for prostate cancer risk. **A)** Violin plot of serum miRs and **B)** serum EV miRs that are predictive of adverse pathology (AP). The lines on the violin plots represent the mean and first and third quartiles. Significant miRs had a p-value < 0.05. **C)** Significant serum and serum EV miRs were included with the CAPRA score in random forest models to predict AP. The area under the curve (AUC) is reported for CAPRA alone (black), CAPRA + serum miRs (red), and CAPRA + serum EV miRs (blue). **D)** Violin plot of serum miRs and **E**) serum EV miRs that are predictive of low-grade and high-grade PCa. Patients stratified in biopsy Gleason grade group 1 were considered low-grade; patients with biopsy Gleason grade group 2 or higher were considered high-grade. The lines on the violin plots represent the mean and first and third quartiles. Significant miRs had a p-value < 0.05. **F)** Significant serum and serum EV miRs were included with the PBCG risk in random forest models to predict low-grade and high-grade PCa. The area under the curve (AUC) is reported for PBCG alone (black), PBCG + serum miRs (red), and PBCG + serum EV miRs (blue).

### Whole serum and serum EV microRNAs significantly differ by biopsy Gleason grade group

Prebiopsy Gleason grade group differentiation was a secondary analysis, as serum samples were collected from patients at least one month post-biopsy. All patients were treatment-naïve; therefore, this analysis was completed under the assumption that the serum and serum EV miR landscape was the same pre- and post-biopsy. PBCG, a clinically utilized prebiopsy risk calculator, was used for comparison to classify high-risk PCa or low-risk PCa by biopsy (15). Random forest models showed that serum miRs and serum EV miRs added to the prognostic power of PBCG in distinguishing between low- and high-grade PCa prebiopsy. 15 serum miRs (**Figure 4D**) and 13 serum EV miRs (**Figure 4E**) differed between patients with low-grade and high-grade PCa. PBCG combined with the significant serum miRs had an AUC of 0.705 (95% CI:0.631-0.783) compared to PBCG risk alone (AUC of 0.600, 95% CI:0.521-0.686) in patients with low-grade and high-grade PCa (p = 0.164) (**Figures 4F and Supplementary Table 6**). PBCG combined with the 13 significant serum EV miRs had an AUC of 0.695 (95% CI:0.589-0.792) compared to PBCG risk alone, which had an AUC of 0.600 (95% CI:0.521-0.686) in patients with low-grade and high-grade PCa (p = 0.080) (**Figure 5C and Supplementary Table 6**). Although there were serum and serum EV miRs that were significantly different in patients with low-grade and high-grade PCa at biopsy, the miRs combined with PBCG did not reach a statistically significant difference in AUCs compared with PBCG alone.

## Discussion

In this study, we report miR profiling of patient-derived prostate cells and tissues and the extracellular vesicles released from these sample types for the first time. Prostate-derived miRs also present in the serum and serum EVs of PCa patients led us to develop a miR panel with prognostic value for aggressive PCa using serum collected after prostate biopsy. We determined that miRs within whole serum and serum EVs added prognostic value to the CAPRA nomogram for adverse pathology and that serum EV miRs were more prognostic than whole serum miRs.

Initial characterization of prostate-derived miRs by small RNA-seq revealed new cell-type specific expression of several miRs. Given the regulatory role of miRs in cell fate^62^, it was unsurprising that the miR profiles of patient-derived prostate samples showed clustering based on cell type (stromal versus epithelial) and sample type (cell versus whole tissue slices). Kumar et al. previously identified prostate stromal- and epithelial-specific miRs using microdissection of prostate tissue ^63^. Both Kumar and our study identified miR-141 as epithelial-specific and that miR-143-3p was highly expressed in stromal samples. We newly identified the miR-199 family as stromal-specific, which has not been previously reported in the prostate. The miR-199 family has been shown to have low expression in prostate tumors ^64^. With the new knowledge that miR-199 is predominantly expressed in the prostate stroma, these reported changes in miR-199a expression in PCa compared to benign prostate tissue may be due to low stromal cells in areas of tumor compared to benign^65^.

The cargo sorting of miRs into EVs was initially thought to be a random process that allowed cells to dispel molecular waste ^66^. However, multiple studies have determined that the sorting process is highly selective, with the secreted EV miRs remaining functional after being transferred to another cell ^67, 68^. In our study, robust intracellular cell type-specific miRs did not always result in high EV miRs. Only 24 of the 107 robustly expressed epithelial intracellular miRs were also highly expressed in epithelial EVs. For the stromal compartment, 50 of the 69 robustly expressed stromal miRs overlapped with stromal EV miRs. These findings support the selective packaging of miRs into EVs, particularly in the epithelial cells, as not all intracellular miRs were found in EVs and not all EV miRs were highly expressed intracellularly ^69^.

Pretreatment serum and serum EV miRs added to the prognostic value of the CAPRA nomogram for the adverse pathology outcome, which was the primary clinical outcome of this study. These new findings improve upon our prior study in which the predictive miRs in the “Serum miR Score” had only provided additional predictive information to patients with low-risk PCa^21^. In this current study, serum and serum EV miRs were significantly different in patients with AP, indicating their potential to provide clinical information to patients with both low- and high-risk PCa before making treatment decisions. Additionally, the relative expression levels of the significantly different miRs in this new panel were both higher and lower in patients with AP, compared to all 14 miRs having higher expression in low-risk PCa in our prior study. A strength of this new study is that we included a racially diverse patient cohort and a full range of Gleason grade groups as opposed to the previous study cohort of only white men and no intermediate-risk patients.

The secondary clinical outcome of this study was to determine the ability of circulating miRs to add prognostic value to a current prebiopsy nomogram, PBCG, which could move the miR-based clinical test earlier in the diagnostic pipeline, potentially reducing biopsies in low-risk patients. One limitation is that the serum samples were collected at least one month post-biopsy, so this analysis was completed assuming that the composition of circulating miRs was the same pre- and post-biopsy. In our study cohort, both serum and serum EV miRs significantly differed in patients with low-versus high-grade PCa; however, they only trended towards adding to the prognostic ability of PBCG (p=0.08). As previously discussed, there is inherent noise in the Gleason grade of prostate biopsies given that thorough sampling of the tumors is not possible. Thus, larger cohorts with prebiopsy serum and post-RP pathology are needed to test biomarkers in this space.

Considerable overlap occurred in the significant miRs between serum and serum EV sample types. Some overlap in miRs was expected because serum EVs are a component of whole serum. Additionally, there was an overlap of prognostic miRs in this study compared to the miRs included in the prognostic “Serum miR Score” from our previous study^21^. Two of the 14 miRs were prognostic for AP in both serum and serum EVs (hsa-let-7a-5p and hsa-miR-451a). In the serum EV model, five additional miRs that were significant for AP overlapped with those included in the Serum miR Score^21^, demonstrating reproducibility of serum miRs over multiple platforms and cohorts. Therefore, this study provides evidence that circulating serum EVs have both overlapping and unique miRs compared to circulating miRs in whole serum and may add additional prognostic information for PCa.

Currently, there are no standardized normalization methods for miRs across biological sample types, with even fewer reports regarding EV miRs. For our serum analysis, there were 10 miRs in the normalization panels, as determined by both geNorm and NormFinder, with seven of the miRs being consistent in both panels. This degree of overlap between normalization methods was consistent with other studies ^70^.

This study was limited by the number of patients who chose to undergo RP treatment and, therefore, the number of samples available for prognostic analysis and model configuration. Of 203 patients, only 81 underwent RP treatment. As our patient cohort was smaller than anticipated, we did not use training and testing subgroups for prognostic model generation. However, we used random forest modeling, which allows for the inclusion of all samples in the model creation due to the bagging feature. Given this limitation, these findings would need to be validated in a prospective cohort of patients with PCa.

Considering the high rates of Gleason grade group discordance between biopsy and RP, the addition of serum/serum EV miR quantitation could be used to guide patient decisions to undergo or forgo radical prostatectomy. With more accurate pretreatment risk stratification, patients with low-risk disease may have more confidence in active surveillance protocols and avoid overtreatment, whereas high-risk patients could be appropriately treated with more aggressive measures. Future studies should assess the clinical significance of these prognostic tests and how they affect treatment recommendations and patient decisions^71^.

## AUTHOR CONTRIBUTIONS

M.L.Z and L.N. conceptualized the study, secured funding, analyzed data and wrote the manuscript. M.L.Z, B.K., M.J.S., A.C.S, and C.L. conducted experiments. T.R.L and M.M.C did informatic and statistical analyses. K.V.N. and M.A. secured patient samples and consent forms. P.H.G. edited manuscript and provided patient sera.

## ACKNOWLEDGMENTS

We thank the patient participants who donated their specimens for this research. Blood samples were acquired with the assistance of Dr. Ruben Sauer and Ms. Patrice King-Lee. BioRender was used to prepare Figure 1A. UIC Research Resource Center Cores, Transmission Electron Microscopy Core, and Genomics Core assisted with data collection.

## REFERENCES

1. Siegel, R. L., Miller, K. D. & Jemal, A. Cancer statistics, 2020. CA Cancer J Clin 70, 7–30 (2020).

2. US Preventive Services Task Force et al. Screening for Prostate Cancer: US Preventive Services Task Force Recommendation Statement. JAMA 319, 1901–1913 (2018).

3. Cancer Facts & Figures 2023. (1930).

4. Epstein, J. I. et al. The 2014 International Society of Urological Pathology (ISUP) Consensus Conference on Gleason Grading of Prostatic Carcinoma: Definition of Grading Patterns and Proposal for a New Grading System. Am J Surg Pathol 40, 244–52 (2016).

5. Epstein, J. I. et al. A Contemporary Prostate Cancer Grading System: A Validated Alternative to the Gleason Score. Eur. Urol. 69, 428–435 (2016).

6. Quintana, L. et al. Gleason Misclassification Rate Is Independent of Number of Biopsy Cores in Systematic Biopsy. Urology 91, 143–149 (2016).

7. Mohler, J. L. et al. Prostate Cancer, Version 2.2019, NCCN Clinical Practice Guidelines in Oncology. J. Natl. Compr. Cancer Netw. JNCCN 17, 479–505 (2019).

8. Aas, K. et al. Increased curative treatment is associated with decreased prostate cancer-specific and overall mortality in senior adults with high-risk prostate cancer; results from a national registry-based cohort study. Cancer Med 9, 6646–6657 (2020).

9. Lei, J. H. et al. Systematic review and meta-analysis of the survival outcomes of first-line treatment options in high-risk prostate cancer. Sci Rep 5, 7713 (2015).

10. Hamdy, F. C. et al. 10-Year Outcomes after Monitoring, Surgery, or Radiotherapy for Localized Prostate Cancer. N Engl J Med 375, 1415–1424 (2016).

11. Sanda, M. G. et al. Quality of life and satisfaction with outcome among prostate-cancer survivors. N Engl J Med 358, 1250–61 (2008).

12. Ankerst, D. P. et al. A Contemporary Prostate Biopsy Risk Calculator Based on Multiple Heterogeneous Cohorts. Eur Urol 74, 197–203 (2018).

13. Kattan, M., Eastham, J. & Wheeler, T. Counseling men with prostate cancer: a nomogram for predicting the presence of small, moderately differentiated, confined tumors. J Urol 170, 1792–1797 (2003).

14. Cooperberg, M. R., Hilton, J. F. & Carroll, P. R. The CAPRA-S score: A straightforward tool for improved prediction of outcomes after radical prostatectomy. Cancer 117, 5039–46 (2011).

15. Cooperberg, M. R. et al. Combined value of validated clinical and genomic risk stratification tools for predicting prostate cancer mortality in a high-risk prostatectomy cohort. Eur Urol 67, 326–33 (2015).

16. Cucchiara, V. et al. Genomic Markers in Prostate Cancer Decision Making. Eur Urol 73, 572– 582 (2018).

17. Salami, S. S., et al. Transcriptomic heterogeneity in multifocal prostate cancer. JCI Insight 3, (2018).

18. Mitchell, P. S. et al. Circulating microRNAs as stable blood-based markers for cancer detection. Proc Natl Acad Sci U A 105, 10513–8 (2008).

19. Cozar, J. et al. The role of miRNAs as biomarkers in prostate cancer. Mutat. Res. Mutat. Res. (2019).

20. Ghafouri-Fard, S., Shoorei, H. & Taheri, M. Role of microRNAs in the development, prognosis and therapeutic response of patients with prostate cancer. Gene 759, 144995 (2020).

21. Mihelich, B. L., Maranville, J. C., Nolley, R., Peehl, D. M. & Nonn, L. Elevated serum microRNA levels associate with absence of high-grade prostate cancer in a retrospective cohort. PLoS One 10, e0124245 (2015).

22. van Niel, G., D’Angelo, G. & Raposo, G. Shedding light on the cell biology of extracellular vesicles. Nat Rev Mol Cell Biol 19, 213–228 (2018).

23. Aalberts, M., Stout, T. A. & Stoorvogel, W. Prostasomes: extracellular vesicles from the prostate. Reproduction 147, R1–14 (2014).

24. Linxweiler, J. & Junker, K. Extracellular vesicles in urological malignancies: an update. Nat Rev Urol 17, 11–27 (2020).

25. Wang, J. et al. Exosomal microRNAs as liquid biopsy biomarkers in prostate cancer. Crit Rev Oncol Hematol 145, 102860 (2020).

26. Saber, S. H. et al. Exosomes are the Driving Force in Preparing the Soil for the Metastatic Seeds: Lessons from the Prostate Cancer. Cells 9, (2020).

27. Richards, Z. et al. Prostate Stroma Increases the Viability and Maintains the Branching Phenotype of Human Prostate Organoids. iScience 12, 304–317 (2019).

28. Théry, C., Amigorena, S., Raposo, G. & Clayton, A. Isolation and characterization of exosomes from cell culture supernatants and biological fluids. Curr Protoc Cell Biol Chapter 3, Unit 3.22 (2006).

29. Martin, M. Cutadapt removes adapter sequences from high-throughput sequencing reads. EMBnet.journal 17, 10–12 (2011).

30. Langmead, B., Trapnell, C., Pop, M. & Salzberg, S. L. Ultrafast and memory-efficient alignment of short DNA sequences to the human genome. Genome Biol. 10, R25 (2009).

31. Team, R. C. R: A language and environment for statistical computing. (2013).

32. Robinson, M. D., McCarthy, D. J. & Smyth, G. K. edgeR: a Bioconductor package for differential expression analysis of digital gene expression data. Bioinformatics 26, 139–40 (2010).

33. Zhao, S., Guo, Y., Sheng, Q. & Shyr, Y. Heatmap3: an improved heatmap package with more powerful and convenient features. BMC Bioinformatics 15, P16 (2014).

34. Wickham, H. ggplot2: Elegant Graphics for Data Analysis. (Springer, 2009). doi:10.1007/978-0-387-98141-3.

35. Sun, W., Reich, B. J., Cai, T. T., Guindani, M. & Schwartzman, A. False Discovery Control in Large-Scale Spatial Multiple Testing. J. R. Stat. Soc. Ser. B Stat. Methodol. 77, 59–83 (2015).

36. Benjamini, Y. & Hochberg, Y. Controlling the False Discovery Rate: A Practical and Powerful Approach to Multiple Testing. J. R. Stat. Soc. Ser. B Methodol. 57, 289–300 (1995).

37. Liaw, A. & Wiener, M. Classification and regression by randomForest. R News 2, 18–22 (2002).

38. Kozminski, M. A. et al. Standardizing the definition of adverse pathology for lower risk men undergoing radical prostatectomy. Urol Oncol 34, 415.e1–6 (2016).

39. Bhagirath, D. et al. microRNA-1246 Is an Exosomal Biomarker for Aggressive Prostate Cancer. Cancer Res 78, 1833–1844 (2018).

40. Li, Z. et al. Exosomal microRNA-141 is upregulated in the serum of prostate cancer patients. Onco Targets Ther 9, 139–48 (2016).

41. Zhang, H. L. et al. An elevated serum miR-141 level in patients with bone-metastatic prostate cancer is correlated with more bone lesions. Asian J Androl 15, 231–5 (2013).

42. Selth, L. A. et al. Circulating microRNAs predict biochemical recurrence in prostate cancer patients. Br J Cancer 109, 641–50 (2013).

43. Lodes, M. J. et al. Detection of cancer with serum miRNAs on an oligonucleotide microarray. PLoS One 4, e6229 (2009).

44. Zidan, H. E., Abdul-Maksoud, R. S., Elsayed, W. S. H. & Desoky, E. A. M. Diagnostic and prognostic value of serum miR-15a and miR-16-1 expression among egyptian patients with prostate cancer. IUBMB Life 70, 437–444 (2018).

45. Cochetti, G. et al. Different levels of serum microRNAs in prostate cancer and benign prostatic hyperplasia: evaluation of potential diagnostic and prognostic role. Onco Targets Ther 9, 7545–7553 (2016).

46. Selth, L. A. et al. Discovery of circulating microRNAs associated with human prostate cancer using a mouse model of disease. Int J Cancer 131, 652–61 (2012).

47. Wang, S. Y. et al. miR-19, miR-345, miR-519c-5p serum levels predict adverse pathology in prostate cancer patients eligible for active surveillance. PLoS One 9, e98597 (2014).

48. Tinay, I. et al. Functional roles and potential clinical application of miRNA-345-5p in prostate cancer. Prostate 78, 927–937 (2018).

49. Cheng, H. H. et al. Circulating microRNA profiling identifies a subset of metastatic prostate cancer patients with evidence of cancer-associated hypoxia. PLoS One 8, e69239 (2013).

50. Haldrup, C. et al. Profiling of circulating microRNAs for prostate cancer biomarker discovery. Drug Deliv Transl Res 4, 19–30 (2014).

51. Brase, J. C. et al. Circulating miRNAs are correlated with tumor progression in prostate cancer. Int J Cancer 128, 608–16 (2011).

52. Wa, Q. et al. miR-204-5p Represses Bone Metastasis via Inactivating NF-κB Signaling in Prostate Cancer. Mol Ther Nucleic Acids 18, 567–579 (2019).

53. Fang, Y. et al. Increased serum levels of miR-214 in patients with PCa with bone metastasis may serve as a potential biomarker by targeting PTEN. Oncol Lett 17, 398–405 (2019).

54. Singh, P. K. et al. Serum microRNA expression patterns that predict early treatment failure in prostate cancer patients. Oncotarget 5, 824–40 (2014).

55. Lyu, J. et al. Discovery and Validation of Serum MicroRNAs as Early Diagnostic Biomarkers for Prostate Cancer in Chinese Population. Biomed Res Int 2019, 9306803 (2019).

56. Kolluru, V. et al. miR-301a expression: Diagnostic and prognostic marker for prostate cancer. Urol Oncol 36, 503.e9–503.e15 (2018).

57. Lieb, V. et al. Serum levels of miR-320 family members are associated with clinical parameters and diagnosis in prostate cancer patients. Oncotarget 9, 10402–10416 (2018).

58. Wach, S. et al. Exploring the MIR143-UPAR Axis for the Inhibition of Human Prostate Cancer Cells In Vitro and In Vivo. Mol Ther Nucleic Acids 16, 272–283 (2019).

59. Mahn, R. et al. Circulating microRNAs (miRNA) in serum of patients with prostate cancer. Urology 77, 1265.e9–16 (2011).

60. Cooperberg, M. R., Broering, J. M. & Carroll, P. R. Risk assessment for prostate cancer metastasis and mortality at the time of diagnosis. J Natl Cancer Inst 101, 878–87 (2009).

61. Cooperberg, M. R., et al. The University of California, San Francisco Cancer of the Prostate Risk Assessment score: a straightforward and reliable preoperative predictor of disease recurrence after radical prostatectomy. J Urol 173, 1938–42 (2005).

62. Park, C. Y., Choi, Y. S. & McManus, M. T. Analysis of microRNA knockouts in mice. Hum. Mol. Genet. 19, R169–R175 (2010).

63. Kumar, B. et al. Cell-type specific expression of oncogenic and tumor suppressive microRNAs in the human prostate and prostate cancer. Sci Rep 8, 7189 (2018).

64. Zhong, J. et al. Downregulation of miR-199a-5p promotes prostate adeno-carcinoma progression through loss of its inhibition of HIF-1alpha. Oncotarget 8, 83523–83538 (2017).

65. Boufaied, N. et al. Development of a predictive model for stromal content in prostate cancer samples to improve signature performance. J Pathol 249, 411–424 (2019).

66. Yuana, Y., Sturk, A. & Nieuwland, R. Extracellular vesicles in physiological and pathological conditions. Blood Rev 27, 31–9 (2013).

67. Melo, S. A. et al. Cancer exosomes perform cell-independent microRNA biogenesis and promote tumorigenesis. Cancer Cell 26, 707–21 (2014).

68. Janas, T., Janas, M. M., Sapoń, K. & Janas, T. Mechanisms of RNA loading into exosomes. FEBS Lett 589, 1391–8 (2015).

69. Liu, X. M., Ma, L. & Schekman, R. Selective sorting of microRNAs into exosomes by phase-separated YBX1 condensates. Elife 10, (2021).

70. Kok, M. G. M. et al. Normalization panels for the reliable quantification of circulating microRNAs by RT-qPCR. FASEB J. Off. Publ. Fed. Am. Soc. Exp. Biol. 29, 3853–3862 (2015).

71. Murphy, A. B. et al. Impact of a Genomic Test on Treatment Decision in a Predominantly African American Population With Favorable-Risk Prostate Cancer: A Randomized Trial. J. Clin. Oncol. Off. J. Am. Soc. Clin. Oncol. 39, 1660–1670 (2021).

