## Supplemental Figures and Tables for "Prostate-derived circulating microRNAs add prognostic value to prostate cancer risk calculators"

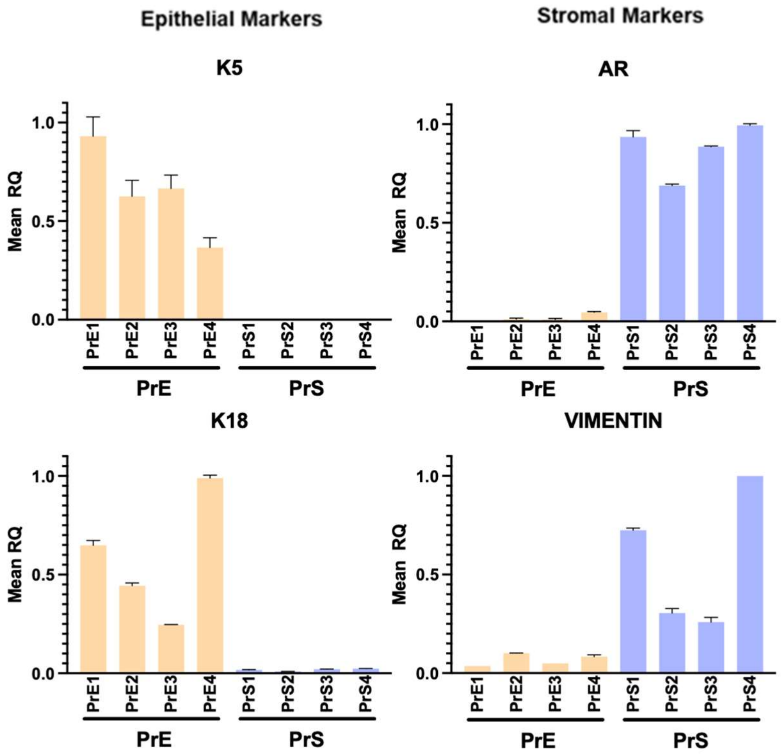


**Supplementary Figure 1. Characterization of patient-derived prostate cells.** RT-qPCR validation of cell-specificity of patient-derived prostate epithelial and stromal cells. Keratin 5 (K5) and Keratin 18 (K18) are epithelial cell markers. Androgen receptor (AR) and Vimentin (VIM) are stromal cell markers. These PrE and PrS samples are the same as those used for the NGS miR profiling studies. (N = 4 patient-derived PrE and PrS, N = 2 replicates per sample, Error bars = SD).

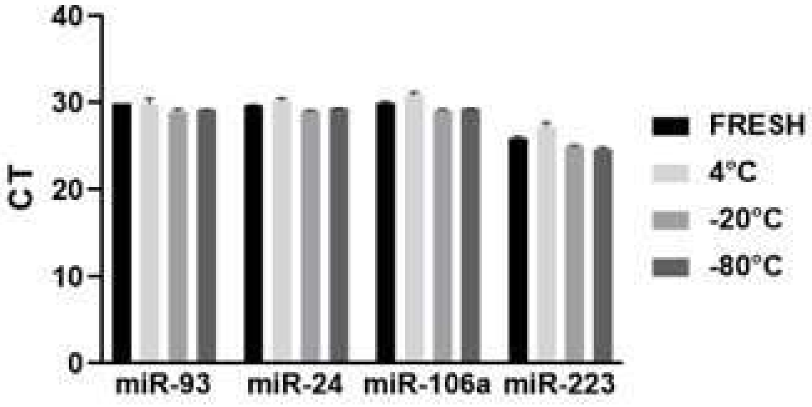


**Supplemental Figure 2.** qPCR testing of miR stability within patient-derived serum EVs stored at 4, -20, or -80°C for 7 days (Black bars = freshly isolated EVs, light/medium/dark gray bars = EVs isolated at 7 days; Error bars = SD).

**Supplementary Table 1. Differentially expressed microRNAs between prostate tissue slices, epithelial cells, and stromal cells.** Differentially expressed microRNAs between prostate tissue slices and prostate epithelial cells, prostate tissue slices and prostate stromal cells, or prostate epithelial cells and prostate stromal cells. The adjusted p-value cutoff was set at ≤ 0.01.

| microRNA | FDR | microRNA | FDR | microRNA | FDR |
|---|---|---|---|---|---|
| hsa-miR-150-5p | 2.14E-12 | hsa-miR-4454 | 9.12E-05 | hsa-miR-8064 | 0.0021 |
| hsa-miR-133a-3p | 2.14E-12 | hsa-miR-424-5p | 9.23E-05 | hsa-miR-6734-5p | 0.0021 |
| hsa-miR-1-3p | 5.49E-12 | hsa-miR-3161 | 9.31E-05 | hsa-miR-182-3p | 0.0021 |
| hsa-miR-363-3p | 2.41E-11 | hsa-miR-483-3p | 9.59E-05 | hsa-miR-197-3p | 0.0022 |
| hsa-miR-375 | 3.06E-11 | hsa-miR-758-3p | 0.0001 | hsa-miR-3201 | 0.0022 |
| hsa-miR-223-3p | 4.34E-11 | hsa-miR-4733-3p | 0.0001 | hsa-miR-3976 | 0.0022 |
| hsa-miR-142-5p | 4.34E-11 | hsa-miR-5591-5p | 0.0001 | hsa-miR-23b-5p | 0.0022 |
| hsa-miR-203b-5p | 4.34E-11 | hsa-miR-802 | 0.0001 | hsa-miR-370-5p | 0.0023 |
| hsa-miR-20b-5p | 4.45E-11 | hsa-miR-4760-5p | 0.0001 | hsa-miR-4312 | 0.0024 |
| hsa-miR-141-5p | 4.45E-11 | hsa-miR-27a-5p | 0.0002 | hsa-miR-181a-3p | 0.0024 |
| hsa-miR-145-3p | 4.93E-11 | hsa-miR-139-3p | 0.0002 | hsa-miR-544a | 0.0024 |
| hsa-miR-205-3p | 5.53E-11 | hsa-miR-224-5p | 0.0002 | hsa-miR-665 | 0.0025 |
| hsa-miR-195-5p | 1.40E-10 | hsa-miR-4724-3p | 0.0002 | hsa-miR-887-3p | 0.0025 |
| hsa-miR-145-5p | 1.40E-10 | hsa-miR-708-3p | 0.0002 | hsa-miR-223-5p | 0.0026 |
| hsa-miR-203a-3p | 2.81E-10 | hsa-miR-4699-3p | 0.0002 | hsa-miR-338-3p | 0.0026 |
| hsa-miR-142-3p | 4.29E-10 | hsa-miR-4484 | 0.0002 | hsa-miR-5088-5p | 0.0027 |
| hsa-miR-199b-5p | 9.51E-10 | hsa-miR-708-5p | 0.0002 | hsa-miR-4438 | 0.0027 |
| hsa-miR-182-5p | 2.18E-09 | hsa-miR-6730-3p | 0.0002 | hsa-miR-34b-3p | 0.0027 |
| hsa-miR-200a-3p | 3.22E-09 | hsa-miR-493-3p | 0.0002 | hsa-miR-1197 | 0.0027 |
| hsa-miR-143-3p | 3.22E-09 | hsa-miR-30d-5p | 0.0002 | hsa-miR-6507-5p | 0.0028 |
| hsa-miR-1-5p | 4.59E-09 | hsa-miR-299-3p | 0.0002 | hsa-miR-374b-5p | 0.0028 |
| hsa-miR-183-5p | 5.16E-09 | hsa-miR-376b-5p | 0.0002 | hsa-miR-140-3p | 0.0028 |
| hsa-miR-135b-5p | 5.16E-09 | hsa-miR-23b-3p | 0.0002 | hsa-miR-29b-2-5p | 0.0029 |
| hsa-miR-96-5p | 5.16E-09 | hsa-miR-320d | 0.0002 | hsa-miR-6512-5p | 0.0029 |
| hsa-miR-99a-5p | 5.77E-09 | hsa-miR-3153 | 0.0002 | hsa-miR-5011-5p | 0.0030 |
| hsa-miR-125b-2-3p | 5.77E-09 | hsa-miR-2355-3p | 0.0003 | hsa-miR-5009-3p | 0.0030 |
| hsa-miR-429 | 6.53E-09 | hsa-miR-133a-5p | 0.0003 | hsa-miR-3928-5p | 0.0030 |
| hsa-miR-200b-3p | 6.53E-09 | hsa-miR-4532 | 0.0003 | hsa-miR-3609 | 0.0030 |
| hsa-miR-7515 | 7.85E-09 | hsa-miR-562 | 0.0003 | hsa-miR-4488 | 0.0030 |
| hsa-miR-143-5p | 7.85E-09 | hsa-miR-8075 | 0.0003 | hsa-miR-877-3p | 0.0033 |
| hsa-miR-214-3p | 1.27E-08 | hsa-miR-664a-3p | 0.0003 | hsa-miR-548v | 0.0033 |
| hsa-miR-504-5p | 1.91E-08 | hsa-miR-4735-5p | 0.0003 | hsa-miR-4481 | 0.0034 |
| hsa-miR-543 | 2.33E-08 | hsa-miR-548aw | 0.0003 | hsa-miR-30e-3p | 0.0034 |
| hsa-miR-126-5p | 2.48E-08 | hsa-miR-192-5p | 0.0003 | hsa-miR-621 | 0.0037 |
| hsa-miR-1293 | 2.61E-08 | hsa-miR-6789-5p | 0.0003 | hsa-miR-33a-5p | 0.0037 |
| hsa-miR-200c-3p | 2.61E-08 | hsa-miR-599 | 0.0003 | hsa-miR-29c-3p | 0.0037 |
| hsa-miR-141-3p | 2.81E-08 | hsa-miR-337-5p | 0.0003 | hsa-miR-2053 | 0.0037 |
| hsa-miR-199a-5p | 2.81E-08 | hsa-miR-616-5p | 0.0004 | hsa-miR-765 | 0.0038 |
| hsa-miR-205-5p | 3.17E-08 | hsa-miR-503-3p | 0.0004 | hsa-miR-6777-5p | 0.0038 |
| hsa-miR-135a-5p | 3.56E-08 | hsa-miR-6735-5p | 0.0004 | hsa-miR-6804-5p | 0.0039 |
| hsa-miR-10b-5p | 3.68E-08 | hsa-miR-28-5p | 0.0004 | hsa-miR-193b-5p | 0.0039 |
| hsa-miR-326 | 3.85E-08 | hsa-miR-5585-3p | 0.0004 | hsa-miR-432-5p | 0.0041 |
| hsa-miR-4483 | 4.56E-08 | hsa-miR-6510-3p | 0.0004 | hsa-miR-1909-3p | 0.0042 |
| hsa-miR-126-3p | 5.09E-08 | hsa-miR-10a-3p | 0.0004 | hsa-miR-30b-5p | 0.0042 |

**Supplementary Table 2. Differentially expressed microRNAs between prostate tissue slice EVs, epithelial cell EVs, and stromal cell EVs.** Differentially expressed microRNAs between prostate tissue slice EVs and prostate epithelial cell EVs, prostate tissue slice EVs and prostate stromal cell EVs, or prostate epithelial cell EVs and prostate stromal cell EVs. The adjusted p-value cutoff was set at ≤ 0.01.

| microRNA | FDR | microRNA | FDR | microRNA | FDR |
|---|---|---|---|---|---|
| hsa-miR-195-5p | 5.79E-07 | hsa-miR-4498 | 0.0002 | hsa-miR-6736-3p | 0.0018 |
| hsa-miR-143-3p | 8.79E-07 | hsa-miR-760 | 0.0002 | hsa-miR-22-5p | 0.0018 |
| hsa-miR-497-5p | 1.32E-06 | hsa-miR-187-3p | 0.0002 | hsa-miR-628-5p | 0.0018 |
| hsa-miR-133a-3p | 1.32E-06 | hsa-let-7a-3p | 0.0002 | hsa-miR-3132 | 0.0018 |
| hsa-miR-143-5p | 1.36E-06 | hsa-miR-99b-3p | 0.0002 | hsa-miR-5194 | 0.0018 |
| hsa-miR-145-5p | 1.36E-06 | hsa-miR-574-3p | 0.0002 | hsa-miR-6130 | 0.0020 |
| hsa-miR-199a-3p | 1.40E-06 | hsa-miR-622 | 0.0002 | hsa-miR-5584-5p | 0.0020 |
| hsa-miR-29c-3p | 1.40E-06 | hsa-let-7a-5p | 0.0002 | hsa-miR-6835-5p | 0.0020 |
| hsa-miR-145-3p | 1.40E-06 | hsa-miR-3131 | 0.0003 | hsa-miR-8063 | 0.0021 |
| hsa-miR-378a-3p | 1.88E-06 | hsa-miR-30b-5p | 0.0003 | hsa-miR-1277-5p | 0.0021 |
| hsa-miR-125b-2-3p | 2.04E-06 | hsa-let-7i-5p | 0.0003 | hsa-miR-3938 | 0.0021 |
| hsa-miR-101-3p | 2.60E-06 | hsa-let-7c-3p | 0.0003 | hsa-miR-141-5p | 0.0022 |
| hsa-miR-199b-3p | 3.95E-06 | hsa-miR-183-5p | 0.0003 | hsa-miR-7515 | 0.0022 |
| hsa-miR-193a-5p | 4.11E-06 | hsa-miR-629-5p | 0.0003 | hsa-miR-532-3p | 0.0022 |
| hsa-miR-16-2-3p | 4.20E-06 | hsa-miR-339-3p | 0.0003 | hsa-miR-3621 | 0.0022 |
| hsa-miR-375 | 4.20E-06 | hsa-miR-3155a | 0.0003 | hsa-miR-136-5p | 0.0022 |
| hsa-miR-30d-5p | 4.51E-06 | hsa-let-7f-1-3p | 0.0003 | hsa-miR-487b-3p | 0.0022 |
| hsa-miR-28-5p | 4.79E-06 | hsa-miR-200b-5p | 0.0003 | hsa-miR-491-5p | 0.0022 |
| hsa-miR-30e-5p | 4.95E-06 | hsa-miR-101-5p | 0.0003 | hsa-miR-4676-3p | 0.0022 |
| hsa-miR-27b-3p | 4.95E-06 | hsa-miR-182-5p | 0.0003 | hsa-miR-597-3p | 0.0022 |
| hsa-miR-23b-3p | 4.95E-06 | hsa-miR-29b-3p | 0.0003 | hsa-miR-345-3p | 0.0023 |
| hsa-miR-141-3p | 5.84E-06 | hsa-miR-374b-5p | 0.0003 | hsa-miR-6809-3p | 0.0023 |
| hsa-miR-21-5p | 6.33E-06 | hsa-miR-628-3p | 0.0003 | hsa-miR-5009-3p | 0.0024 |
| hsa-let-7c-5p | 6.33E-06 | hsa-miR-194-5p | 0.0003 | hsa-miR-6874-5p | 0.0024 |
| hsa-miR-99a-5p | 6.95E-06 | hsa-miR-151a-3p | 0.0004 | hsa-miR-5690 | 0.0024 |
| hsa-miR-328-3p | 7.64E-06 | hsa-miR-4532 | 0.0004 | hsa-miR-181c-5p | 0.0024 |
| hsa-miR-26a-5p | 7.93E-06 | hsa-miR-381-3p | 0.0004 | hsa-miR-1243 | 0.0024 |
| hsa-miR-28-3p | 8.65E-06 | hsa-miR-562 | 0.0004 | hsa-miR-154-5p | 0.0024 |
| hsa-miR-140-3p | 8.74E-06 | hsa-miR-98-5p | 0.0004 | hsa-miR-376a-3p | 0.0025 |
| hsa-miR-20b-5p | 8.74E-06 | hsa-miR-378g | 0.0004 | hsa-miR-3682-3p | 0.0025 |
| hsa-miR-1-3p | 8.75E-06 | hsa-miR-660-3p | 0.0004 | hsa-miR-412-5p | 0.0025 |
| hsa-miR-146a-5p | 9.61E-06 | hsa-miR-125b-1-3p | 0.0004 | hsa-miR-3126-3p | 0.0026 |
| hsa-miR-23b-5p | 1.03E-05 | hsa-miR-320d | 0.0004 | hsa-miR-4483 | 0.0026 |
| hsa-miR-214-3p | 1.06E-05 | hsa-miR-2681-3p | 0.0004 | hsa-miR-3144-3p | 0.0026 |
| hsa-miR-125a-5p | 1.06E-05 | hsa-miR-127-3p | 0.0004 | hsa-miR-411-5p | 0.0026 |
| hsa-miR-148b-3p | 1.08E-05 | hsa-miR-7977 | 0.0004 | hsa-miR-6865-5p | 0.0027 |
| hsa-miR-22-3p | 1.13E-05 | hsa-miR-574-5p | 0.0004 | hsa-miR-487b-5p | 0.0028 |
| hsa-miR-365a-3p | 1.18E-05 | hsa-miR-335-5p | 0.0004 | hsa-miR-1307-3p | 0.0028 |
| hsa-miR-221-3p | 1.18E-05 | hsa-miR-132-3p | 0.0004 | hsa-miR-4459 | 0.0028 |
| hsa-miR-484 | 1.18E-05 | hsa-miR-103a-3p | 0.0004 | hsa-let-7e-3p | 0.0028 |
| hsa-miR-532-5p | 1.18E-05 | hsa-miR-423-5p | 0.0004 | hsa-miR-1260a | 0.0028 |
| hsa-miR-378c | 1.18E-05 | hsa-miR-4699-3p | 0.0004 | hsa-miR-4735-5p | 0.0028 |
| hsa-miR-152-3p | 1.18E-05 | hsa-miR-3135b | 0.0004 | hsa-miR-4328 | 0.0029 |
| hsa-miR-130a-3p | 1.25E-05 | hsa-miR-6735-5p | 0.0004 | hsa-miR-7706 | 0.0030 |

**Supplementary Table 3. MicroRNAs specific to epithelial cell, stromal cell, tissue slice explant, and overlapping groups.** TMM-normalized miRs with > 500 counts per million (cpm) in patient-derived epithelial cells (PrE), stromal cells (PrS), and tissue slice explants (TSC).

| Sample Group | miRs | | | |
|---|---|---|---|---|
| PrE/PrS/TSC | hsa-miR-29c-3p | hsa-miR-15b-5p | hsa-miR-25-3p | hsa-let-7f-5p |
| | hsa-miR-103a-3p | hsa-let-7d-5p | hsa-miR-98-5p | hsa-miR-152-3p |
| | hsa-miR-27a-3p | hsa-miR-22-3p | hsa-miR-127-3p | hsa-miR-30e-5p |
| | hsa-miR-93-5p | hsa-miR-27b-3p | hsa-let-7b-5p | hsa-miR-101-3p |
| | hsa-let-7c-5p | hsa-miR-29b-3p | hsa-miR-34a-5p | hsa-miR-30a-5p |
| | hsa-miR-128-3p | hsa-miR-30d-5p | hsa-miR-100-5p | hsa-miR-222-3p |
| | hsa-miR-151a-3p | hsa-miR-30c-5p | hsa-let-7i-5p | hsa-miR-148b-3p |
| | hsa-miR-29a-3p | hsa-let-7a-5p | hsa-miR-148a-3p | hsa-miR-26b-5p |
| | hsa-let-7g-5p | hsa-miR-143-3p | hsa-miR-130a-3p | hsa-miR-191-5p |
| | hsa-miR-16-5p | hsa-miR-99b-5p | hsa-miR-23b-3p | hsa-miR-218-5p |
| | hsa-miR-24-3p | hsa-miR-221-3p | hsa-miR-26a-5p | hsa-miR-125b-5p |
| | hsa-miR-23a-3p | hsa-miR-186-5p | hsa-miR-125a-5p | hsa-miR-21-5p |
| | hsa-miR-19b-3p | hsa-miR-224-5p | hsa-let-7e-5p | |
| PrE/PrS | hsa-miR-7-5p | | | |
| | hsa-miR-210-3p | | | |
| | hsa-miR-31-5p | | | |
| | hsa-miR-424-5p | | | |
| PrE/TSC | hsa-miR-320a | hsa-miR-200a-3p | hsa-miR-30b-5p | hsa-miR-361-3p |
| | hsa-miR-374a-5p | hsa-miR-28-3p | hsa-miR-20a-5p | hsa-miR-183-5p |
| | hsa-miR-126-3p | hsa-miR-361-5p | hsa-miR-182-5p | hsa-miR-30a-3p |
| | hsa-miR- | hsa-miR-205-5p | hsa-miR-203a-3p | hsa-miR-92a-3p |
| | 151b/151a-5p | hsa-miR-15a-5p | hsa-miR-454-3p | hsa-miR-200c-3p |
| | hsa-miR-141-3p | hsa-miR-423-5p | hsa-miR-149-5p | hsa-miR-200b-3p |
| | hsa-miR-429 | hsa-miR-342-3p | hsa-miR-365b-3p | hsa-miR-335-5p |
| | hsa-miR-365a-3p | hsa-miR-339-5p | | |
| | hsa-miR-107 | | | |
| PrS/TSC | hsa-miR-146a-5p | | | |
| | hsa-miR-199a-3p | | | |
| | hsa-miR-155-5p | | | |
| | hsa-miR-145-5p | | | |
| | hsa-miR-199a-5p | | | |
| | hsa-miR-574-3p | | | |
| | hsa-miR-199b-3p | | | |
| PrE | hsa-miR-141-5p | hsa-miR-103b | hsa-miR-301a-3p | hsa-miR-4784 |
| | hsa-miR-192-5p | hsa-miR-185-5p | hsa-miR-19a-3p | hsa-miR-590-3p |
| | hsa-miR-135b-5p | hsa-miR-31-3p | hsa-miR-542-3p | hsa-miR-96-5p |
| | hsa-let-7a-3p | hsa-miR-503-5p | hsa-miR-138-5p | hsa-miR-181b-5p |
| | hsa-miR-21-3p | hsa-miR-455-5p | hsa-miR-708-5p | hsa-miR-302b-5p |
| | hsa-miR-203b-5p | hsa-miR-17-5p | | |
| PrS | hsa-miR-379-5p | | | |
| | hsa-miR-134-5p | | | |
| | hsa-miR-376a-3p | | | |
| | hsa-miR-409-3p | | | |
| | hsa-miR-432-5p | | | |
| | hsa-miR-382-5p | | | |
| | hsa-miR-574-5p | | | |
| TSC | hsa-miR-375 | hsa-miR-6865-5p | hsa-miR-20b-5p | hsa-miR-223-3p |
| | hsa-miR-150-5p | hsa-miR-142-3p | hsa-miR-363-3p | hsa-miR-214-3p |
| | hsa-miR-29c-5p | hsa-miR-497-5p | hsa-miR-133a-3p | hsa-miR-145-3p |
| | hsa-miR-142-5p | hsa-miR-532-5p | hsa-miR-135a-5p | hsa-miR-10a-5p |
| | hsa-miR-143-5p | hsa-miR-1-3p | hsa-miR-199b-5p | hsa-miR-374b-5p |
| | hsa-miR-146b-5p | hsa-miR-10b-5p | hsa-miR-140-3p | hsa-miR-195-5p |
| | hsa-miR-196b-5p | hsa-miR-99a-5p | hsa-miR-378a-3p | hsa-miR-30e-3p |
| | hsa-miR-126-5p | | | |

**Supplementary Table 4. MicroRNAs specific to epithelial cell EVs, stromal cell EVs, tissue slice explant EVs, serum EVs, and overlapping groups**

| Sample Group | miRs | | | |
|---|---|---|---|---|
| PrE EV/PrS EV/TSC EV/Serum EV | hsa-miR-29a-3p | hsa-let-7a-5p | hsa-let-7i-5p | hsa-miR-30a-5p |
| | hsa-miR-16-5p | hsa-miR-143-3p | hsa-miR-26a-5p | hsa-miR-26b-5p |
| | hsa-miR-24-3p | hsa-miR-221-3p | hsa-miR-125a-5p | hsa-miR-125b-5p |
| | hsa-miR-23a-3p | hsa-let-7b-5p | hsa-let-7f-5p | hsa-miR-21-5p |
| | hsa-miR-30d-5p | hsa-miR-148a-3p | | |
| PrE EV/PrS EV/TSC EV | hsa-miR-29b-3p | | | |
| | hsa-miR-31-5p | | | |
| PrE EV/ TSC EV/ Serum EV | None | | | |
| PrS EV/TSC EV/Serum EV | hsa-miR-103a-3p | hsa-miR-146a-5p | hsa-miR-423-5p | hsa-miR-140-3p |
| | hsa-miR-320a | hsa-miR-122-5p | hsa-miR-155-5p | hsa-miR-223-3p |
| | hsa-miR-126-3p | hsa-miR-199a-3p | hsa-miR-342-3p | hsa-miR-30e-5p |
| | hsa-miR-27a-3p | hsa-let-7g-5p | hsa-miR-25-3p | hsa-miR-148b-3p |
| | hsa-miR-93-5p | hsa-miR-361-5p | hsa-miR-339-5p | hsa-miR-92a-3p |
| | hsa-miR-128-3p | hsa-miR-22-3p | hsa-miR-23b-3p | hsa-miR-191-5p |
| | hsa-miR-151a-3p | hsa-miR-142-3p | hsa-let-7e-5p | hsa-miR-486-5p |
| | hsa-let-7c-5p | hsa-miR-27b-3p | | |
| PrE EV/TSC EV | hsa-miR-7-5p | | | |
| | hsa-miR-205-5p | | | |
| | hsa-miR-203a-3p | | | |
| | hsa-miR-4301 | | | |
| | hsa-miR-200c-3p | | | |
| PrS EV/TSC EV | hsa-miR-379-5p | hsa-miR-145-5p | hsa-miR-34a-5p | hsa-miR-222-3p |
| | hsa-miR-134-5p | hsa-miR-432-5p | hsa-miR-100-5p | hsa-miR-382-5p |
| | hsa-miR-196b-5p | hsa-miR-127-3p | hsa-miR-152-3p | hsa-miR-199b-3p |
| | hsa-miR-409-3p | | | |
| TSC EV/Serum EV | hsa-miR-29c-3p | hsa-miR-107 | hsa-miR-30c-5p | hsa-miR-425-5p |
| | hsa-miR-192-5p | hsa-miR-185-5p | hsa-miR-186-5p | hsa-miR-361-3p |
| | hsa-miR-150-5p | hsa-miR-106b-3p | hsa-miR-10b-5p | hsa-miR-194-5p |
| | hsa-miR-423-3p | hsa-miR-28-3p | hsa-miR-181a-5p | hsa-let-7d-3p |
| | hsa-miR-142-5p | hsa-miR-15b-5p | hsa-miR-197-3p | hsa-miR-10a-5p |
| | hsa-miR-744-5p | hsa-miR-19b-3p | hsa-miR-20a-5p | hsa-miR-484 |
| | hsa-miR-146b-5p | hsa-let-7d-5p | hsa-miR-182-5p | hsa-miR-1307-3p |
| | hsa-miR-126-5p | hsa-miR-15a-5p | hsa-miR-101-3p | |
| PrE EV | None | | | |
| PrS EV | hsa-miR-6735-5p | hsa-miR-1181 | hsa-miR-4532 | hsa-miR-6748-5p |
| | hsa-miR-543 | hsa-miR-3940-3p | hsa-miR-211-3p | hsa-miR-6510-3p |
| | hsa-miR-4520-3p | hsa-miR-6756-3p | hsa-miR-8075 | hsa-miR-5684 |
| | hsa-miR-4498 | hsa-miR-629-3p | hsa-miR-3605-3p | hsa-miR-3131 |
| | hsa-miR-660-3p | hsa-miR-4699-3p | hsa-miR-3607-5p | hsa-miR-302b-5p |
| | hsa-miR-5010-5p | hsa-miR-7706 | | |
| TSC EV | hsa-miR-6126 | hsa-miR-23b-5p | hsa-miR-494-3p | hsa-miR-378a-3p |
| | hsa-miR-375 | hsa-miR-339-3p | hsa-miR-4454 | hsa-miR-365b-3p |
| | hsa-miR-9-5p | hsa-miR-664a-5p | hsa-miR-140-5p | hsa-miR-424-5p |
| | hsa-miR-141-5p | hsa-miR-32-5p | hsa-miR-125b-2-3p | hsa-miR-214-3p |
| | hsa-miR-421 | hsa-let-7b-3p | hsa-miR-363-3p | hsa-miR-125b-1-3p |
| | hsa-miR-374a-5p | hsa-miR-4324 | hsa-miR-210-3p | hsa-miR-92b-3p |
| | hsa-miR-151b/151a-5p | hsa-miR-500a-3p | hsa-miR-200b-5p | hsa-miR-183-5p |
| | hsa-miR-29c-5p | hsa-miR-497-5p | hsa-miR-1247-5p | hsa-miR-145-3p |
| | hsa-miR-802 | hsa-miR-340-5p | hsa-miR-132-3p | hsa-miR-30a-3p |
| | hsa-miR-654-3p | hsa-miR-532-5p | hsa-miR-760 | hsa-miR-378c |
| | hsa-miR-143-5p | hsa-miR-1-3p | hsa-miR-30b-5p | hsa-miR-187-3p |
| | hsa-miR-328-3p | hsa-miR-193b-5p | hsa-miR-130a-3p | hsa-miR-200b-3p |
| | hsa-miR-1307-5p | hsa-miR-324-5p | hsa-miR-324-3p | hsa-miR-574-3p |
| | hsa-miR-133b | hsa-miR-16-2-3p | hsa-miR-629-5p | hsa-miR-335-5p |
| | hsa-let-7a-3p | hsa-miR-99b-5p | hsa-miR-199a-5p | hsa-miR-96-5p |
| | hsa-miR-141-3p | hsa-miR-320b | hsa-miR-381-3p | hsa-miR-574-5p |
| | hsa-miR-320c | hsa-miR-224-5p | hsa-miR-133a-3p | hsa-miR-1246 |
| | hsa-miR-193a-5p | hsa-miR-99a-5p | hsa-miR-345-5p | hsa-miR-99a-3p |
| | hsa-miR-429 | hsa-miR-19a-3p | hsa-miR-221-5p | hsa-miR-218-5p |
| | hsa-miR-365a-3p | hsa-miR-425-3p | hsa-miR-452-5p | hsa-miR-195-5p |
| | hsa-miR-103b | hsa-miR-98-5p | hsa-miR-199b-5p | hsa-miR-1296-5p |
| | hsa-miR-200a-3p | hsa-miR-20b-5p | hsa-miR-149-5p | hsa-miR-130b-3p |
| | hsa-miR-376a-3p | hsa-miR-660-5p | | |
| Serum EV | hsa-miR-584-5p | | | |
| | hsa-miR-223-5p | | | |
| | hsa-miR-486-3p | | | |
| | hsa-miR-3615 | | | |
| | hsa-miR-625-3p | | | |
| | hsa-miR-451a | | | |
| | hsa-miR-4433b-5p | | | |

**Supplementary Table 5. MicroRNAs included in the random forest models with CAPRA for predicting adverse pathology .** The AUC and 95% confidence interval (CI) for CAPRA Score alone is shown the AP outcome. The AUCs and 95% CI for significant serum and serum EV miRs added to CAPRA Score for the AP outcome are also represented. The significant miRs that were included in each model are listed. AUC values were generated with leave-one-out cross validation (LOOCV).

| Model | LOOCV AUC | 95% CI | Variables |
|---|---|---|---|
| CAPRA Score → AP | 0.516 | 0.379-0.643 | CAPRA Score only |
| CAPRA Score + serum miRs → AP | 0.675 | 0.547-0.799 | hsa-let-7a-5p<br>hsa-miR-451a<br>hsa-miR-16-5p<br>hsa-miR-363-3p<br>hsa-miR-30a-5p<br>hsa-miR-122-5p<br>hsa-miR-25-3p<br>CAPRA Score |
| CAPRA Score + serum EV miRs → AP | 0.739 | 0.580-0.900 | hsa-let-7a-5p<br>hsa-miR-103a-3p<br>hsa-miR-107<br>hsa-miR-26b-5p<br>hsa-miR-30c-5p<br>hsa-miR-451a<br>hsa-miR-93-5p<br>hsa-miR-19b-3p<br>hsa-miR-222-3p<br>hsa-miR-16-5p<br>hsa-miR-301a-3p<br>hsa-miR-320a<br>hsa-miR-345-5p<br>hsa-miR-363-3p<br>hsa-miR-191-5p<br>hsa-miR-199a-5p<br>hsa-miR-221-3p<br>hsa-miR-27b-3p<br>hsa-miR-25-3p<br>CAPRA Score |

**Supplementary Table 6. MicroRNAs included in the random forest models with PBCG for predicting low-grade versus high-grade PCa prebiopsy.** The AUCs with 95% confidence intervals for PBCG risk alone is shown for low-grade versus high-grade PCa at biopsy. The AUCs and 95% CI for significant serum and serum EV miRs added to PBCG risk for this outcome are also represented. The significant miRs that were included in each model are listed. AUC values were generated with leave-one-out cross validation (LOOCV). PBCG HG is the high-grade percentage, and PBCG LG is the low-grade percentage, as calculated by PBCG.

| Model | LOOCV AUC | 95% CI | Variables | |
|---|---|---|---|---|
| PBCG Risk → Low-grade/High-grade PCa | 0.600 | 0.521-0.686 | PBCG HG, LG, Negative Risk | |
| PBCG Risk + serum miRs → Low-grade PCa | 0.705 | 0.630-0.783 | hsa-miR-16-5p<br>hsa-miR-19b-3p<br>hsa-let-7b-5p<br>hsa-miR-363-3p<br>hsa-miR-451a<br>hsa-miR-1246<br>hsa-miR-320a<br>hsa-miR-27a-3p<br>hsa-miR-221-3p | hsa-miR-27b-3p<br>hsa-miR-106a-5p<br>hsa-miR-30a-5p<br>hsa-miR-874-3p<br>hsa-miR-23a-3p<br>hsa-miR-191-5p<br>PBCG HG<br>PBCG LG<br>PBCG Negative |
| PBCG Risk + serum EV miRs → Low-grade PCa | 0.695 | 0.589-0.792 | hsa-miR-191-5p<br>hsa-miR-335-5p<br>hsa-miR-363-3p<br>hsa-miR-221-3p<br>hsa-miR-330-3p<br>hsa-miR-19b-3p<br>hsa-miR-103a-3p<br>hsa-miR-34a-5p | hsa-miR-107<br>hsa-miR-130b-3p<br>hsa-miR-16-5p<br>hsa-let-7a-5p<br>hsa-miR-451a<br>PBCG HG<br>PBCG LG<br>PBCG Negative |
